# DNMT1 inhibition by pUG-fold quadruplex RNA

**DOI:** 10.1101/2022.10.14.512272

**Authors:** Linnea I. Jansson-Fritzberg, Camila I. Sousa, Michael J. Smallegan, Jessica J. Song, Anne R. Gooding, Vignesh Kasinath, John L. Rinn, Thomas R. Cech

**Affiliations:** BioFrontiers Institute, University of Colorado Boulder, Boulder, CO, 80303, USA; Department of Biochemistry, University of Colorado Boulder, Boulder, CO, 80303, USA; Department of Molecular, Cellular and Developmental Biology, University of Colorado Boulder, Boulder, CO, 80303, USA; Howard Hughes Medical Institute, University of Colorado Boulder, Boulder, CO, 80303, USA

**Keywords:** RNA, pUG-fold, DNMT1, Epigenetics, G-Quadruplex, DNA methylation

## Abstract

Aberrant DNA methylation is one of the earliest hallmarks of cancer. DNMT1 is responsible for methylating newly replicated DNA, but the precise regulation of DNMT1 to ensure faithful DNA methylation remains poorly understood. A link between RNA and chromatin-associated proteins has recently emerged, and several studies have shown that DNMT1 can be regulated by a variety of RNAs. In this study we have confirmed that human DNMT1 indeed interacts with multiple RNAs, including its own nuclear mRNA. Unexpectedly, we found that DNMT1 exhibits a strong and specific affinity for GU-rich RNAs that form a pUG-fold, a non-canonical G-quadruplex. We find that pUG-fold-capable RNAs inhibit DNMT1 activity by inhibiting binding of hemimethylated DNA, and additionally provide evidence for multiple RNA binding modes with DNMT1. Together, our data indicates that a human chromatin-associated protein binds to and is regulated by pUG-fold RNA.

## INTRODUCTION

Many chromatin-modifying enzymes that bind DNA and histones have also been found to bind RNA. This RNA binding has been proposed to play a role in both the recruitment and regulation of enzyme activity, but detailed mechanisms of RNA-mediated regulation remain poorly understood (Beltran et al. 2019; Davidovich and Cech 2015; Winkler and Dimitrova 2022). Examples of chromatin-associated proteins that exhibit both DNA and RNA binding include PRC2, WDR5, CTCF, RNA polymerase II, many transcription factors, and the DNMT family of methyltransferases (Di Ruscio et al. 2013; Long et al. 2017, 2020; Merry et al. 2015; Ponicsan et al. 2013; Saldaña-Meyer et al. 2019; Wang et al. 2015, 2017).

The DNA methyltransferases methylate cytosines within CpG dinucleotides – a major mechanism of epigenetic gene repression (Bird 2002; Lyko 2018). CpG islands are prevalent near or in many promoters, thus the need for faithful methylation maintenance in these regions is critical (Esteller 2002). The failure to maintain the methylation state of the genome results in genomic instability and disease susceptibility (Baylin and Herman 2000; Eden et al. 2003; Feinberg and Vogelstein 1983). There are three classes of DNMTs; DNMT1 and DNMT3A/B are DNA methyltransferases while DNMT2 methylates tRNAs (Goll et al. 2006; Goll and Bestor 2005; Jeltsch et al. 2016; Lyko 2018). DNMT1 is a maintenance DNA methyltransferase that restores the methylation state to newly replicated hemimethylated DNA, and DNMT3A/B are de novo methyltransferases that deposit methyl marks during development and differentiation (Gao et al. 2020; Gruenbaum, Cedar, and Razin 1982; Jeltsch 2006; W. Ren, Gao, and Song 2018). While these distinctions are generally true, there has been evidence of de novo activity by DNMT1 as well (Haggerty et al. 2021; Wang et al. 2020). DNMT1 has been shown to interact with, and be regulated by, a variety of cofactors that include other proteins, small molecules and RNA (Di Ruscio et al. 2013; Estève et al. 2006; G Hendrickson et al. 2016; Merry et al. 2015; R. Ren et al. 2018; W. Ren et al. 2021).

DNMT1 has been demonstrated to directly associate with a variety of RNAs, which seem to have varying effects on DNMT1 activity and localization. DACOR1, a long non-coding RNA (lncRNA) which is downregulated in colon cancer, was shown to interact with DNMT1 (Merry et al. 2015). Loss of this interaction promoted global hypomethylation, although the precise mode of interaction between this RNA and DNMT1 remains unclear (Merry et al. 2015; Somasundaram et al. 2018). Additionally, DNMT1 has been shown to directly interact with the lncRNA ecCEBPA, an antisense lncRNA originating from the *CEBPA* locus (Di Ruscio et al. 2013). DNMT1 is a large protein (183 kDa) with a large regulatory N-terminal region and a catalytic C-terminal domain. ecCEBPA RNA was shown to bind to the C-terminal region of DNMT1 and promote local hypomethylation at its native locus by preventing DNMT1 from binding the promoter, thereby allowing for transcription of the *CEBPA* gene. In the same study, genome-wide analysis indicated that inhibition of DNMT1 action by nascent RNA was general beyond the *CEBPA* locus.

Additionally, DNMT1 has been shown to be inhibited by binding to various miRNAs, including miR-155-5p (Zhang et al. 2015). This RNA interacts with the C-terminal region of the protein and was shown to be inhibitory to DNA binding and therefore methyltransferase activity. This inhibition was seen with various RNA constructs, including RNAs with G-quadruplex-forming potential.

In this study, we investigate the sequence and structure specificity of DNMT1-RNA interactions and the effect of RNA binding on methyltransferase activity. We first use formaldehyde RNA immunoprecipitation followed by next-generation sequencing (fRIP-seq) on endogenous DNMT1 in two different cell lines and find that DNMT1 interacts with many RNAs, and surprisingly, its own mRNA. Upon further analysis of RNA sequence and structure preference of DNMT1, we see a high and specific affinity for a recently discovered RNA structure called a poly-(UG)-fold (pUG-fold) (Roschdi et al. 2021). We also find that DNMT1 displays a general but lower affinity for a variety of other RNAs. Finally, we demonstrate that binding of pUG-fold RNA to DNMT1 inhibits DNMT1 methyltransferase activity.

## RESULTS

### DNMT1 binds to many different RNAs in the nucleus, including its own mRNA

To dissect the binding preference of DNMT1 to various RNAs we performed DNMT1 fRIP-seq in two different human cell lines, the leukemia cell line K562 and induced Pluripotent Stem Cells (iPSCs) (**Fig 1A, Supplemental Fig S1A, B**). Additionally, we performed fRIP-seq with two different DNMT1-targeting antibodies to identify consistent RNA binding events. There was good overlap between the datasets from both antibody pulldowns in both cell types (**Supplemental Fig S1C, D**). DNMT1 bound about 3300 different RNAs in K562 cells and about 1200 in iPSCs (Log2FoldChange > 1, Adjusted p-value (padj) < 0.01); this difference correlates with the higher level of DNMT1 in the K562 cell line as measured by western blot signal intensity **(Fig 1A, Supplemental Fig S1A, B)**. One of the most significantly enriched RNAs, regardless of cell line or antibody, was the *DNMT1* mRNA (**Fig 1A, Supplemental Fig S1B**).

**FIGURE 1.**
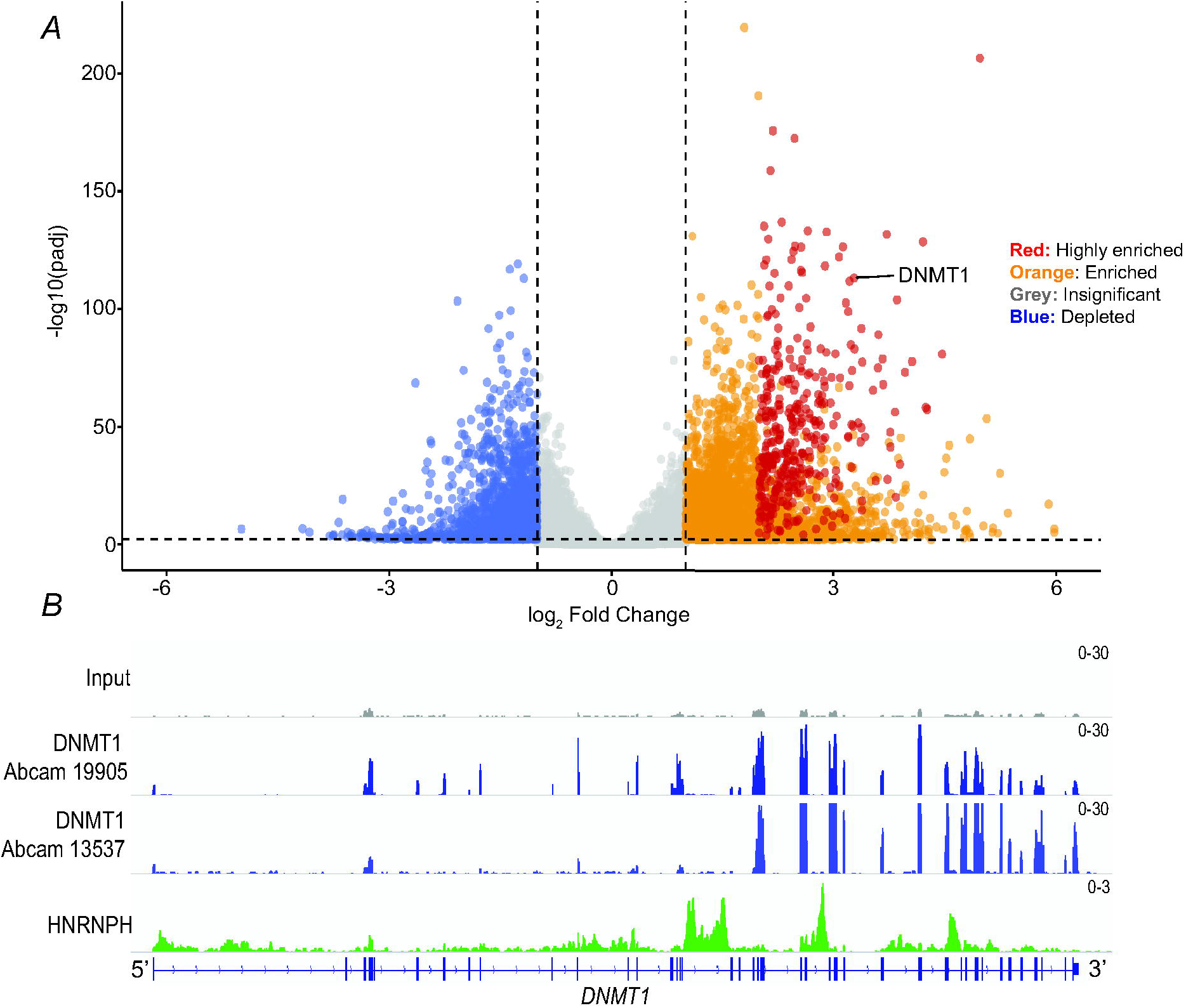
DNMT1 has numerous binding partners including its own mRNA. (*A*) A volcano plot showing enriched RNAs from fRIP-seq in K562 cells. Highly enriched RNAs (red) have been classified by a Log_2_FoldChange > 2, baseMean > 100 and adjusted p-value (padj) < 0.001. Enriched RNAs (orange) are classified by a Log_2_FoldChange > 1, baseMean > 100 and padj < 0.001. Non-significant and depleted RNAs are shown in grey and blue, respectively. (*B*) Integrated Genomics Viewer (IGV) track of DNMT1 (blue) and HnRNP H (green) fRIP peaks over the *DNMT1* mRNA.

The highest percentage of DNMT1 fRIP-seq peaks were within introns and regulatory regions (**Supplemental Fig S2A**). However, the interaction between DNMT1 and its mRNA was biased to the fully spliced and processed mRNA as evidenced by peaks observed mainly over exonic regions (**Fig 1B**). To check if our fRIP-seq assay was biasing the observed results, we used HnRNP H, a known intronic binding protein, as a control and saw peaks mainly within introns of the same mRNA (**Fig 1B, Supplemental Fig S2A**). Because DNMT1 is a nuclear protein, its preference for binding to the fully spliced transcript was unexpected. To test if the observed interaction between DNMT1 and its mRNA could be due to crosslinking of the nascent DNMT1 peptide and the mRNA in the cytoplasm during translation, we performed the same fRIP-seq experiments after nuclear fractionation (**Supplemental Fig S2B**). We observed the same DNMT1-mRNA interaction in the nucleus, thereby supporting the hypothesis that this interaction is a nuclear binding event (**Supplemental Fig S2C**). In addition to binding to the fully spliced transcript, we noticed a preference for binding towards the 3’ end of the mRNA. Although there was a slight PCR amplification bias towards the 3’ end of the *DNMT1* mRNA, it was not enough to explain the observed 3’-binding preference of DNMT1 (**Supplemental Fig S2D**).

### DNMT1 shows binding preference for GU-repeat containing RNA

To further test the affinity of DNMT1 to its own mRNA, we made six ∼200 nt RNA truncations by in vitro transcription (**Fig 2A, Supplemental Table 1**). These RNAs span the regions that show peaks in the fRIP-seq of Fig 1B (**Fig 2A**). To analyze binding of these RNAs, we performed electrophoretic mobility shift assays (EMSAs) with recombinant full-length human DNMT1 (**Supplemental Fig S3A**). While all RNAs bound with similar affinities, RNA3 had the highest affinity (∼250 nM *K*_*d*_^*app*^) (**Fig 2B, D, Table 1**). Several of the RNAs (# 7,12,13,25) had broad binding curves, which we interpret to indicate that they fold into two or more conformers that have different affinities for DNMT1. This notion is further supported by the presence of multiple bands for these RNAs when run on a native gel (**Supplemental Fig S3B**). In general, the affinity of DNMT1 for the various mRNA truncations is similar regardless of sequence composition or predicted amount of secondary structure, indicating that DNMT1 has a broad general affinity for RNA.

**TABLE 1.** Equilibrium constants for binding of various RNAs to DNMT1

| RNA/DNA | Length (nt) | $K_d^{app}$ (nM) in KCl | Hill coeff in KCl | $K_d^{app}$ (nM) in LiCl | Hill coeff in LiCl | Fold Change $K_d^{app}$ in LiCl/KCl |
| --- | --- | --- | --- | --- | --- | --- |
| **RNA3** | 208 | $262 \pm 3$ | $2.34 \pm 0.05$ | N/A | N/A | N/A |
| **RNA7** | 253 | $282 \pm 13$ | $0.75 \pm 0.03$ | N/A | N/A | N/A |
| **RNA12** | 225 | $391 \pm 13$ | $0.88 \pm 0.03$ | N/A | N/A | N/A |
| **RNA13** | 222 | $482 \pm 52$ | $0.52 \pm 0.03$ | N/A | N/A | N/A |
| **RNA23** | 205 | $332 \pm 6$ | $1.31 \pm 0.03$ | N/A | N/A | N/A |
| **RNA25** | 200 | $371 \pm 25$ | $0.71 \pm 0.04$ | N/A | N/A | N/A |
| **(C)<sub>40</sub>** | 40 | $390 \pm 8$ | $1.45 \pm 0.04$ | N/A | N/A | N/A |
| **(G)<sub>40</sub>** | 40 | $1030 \pm 13$ | $2.44 \pm 0.07$ | N/A | N/A | N/A |
| **(U)<sub>40</sub>** | 40 | $1711 \pm 57$ | $1.01 \pm 0.04$ | N/A | N/A | N/A |
| **(A)<sub>40</sub>** | 40 | $5348 \pm 743$ | $5.8 \pm 2.77$ | N/A | N/A | N/A |
| **(GA)<sub>20</sub>** | 40 | $348 \pm 9$ | $1.41 \pm 0.04$ | N/A | N/A | N/A |
| **(GGAA)<sub>10</sub>** | 40 | $216 \pm 10$ | $0.98 \pm 0.04$ | N/A | N/A | N/A |
| **(CGG)<sub>12</sub>** | 36 | $1074 \pm 21$ | $1.9 \pm 0.06$ | N/A | N/A | N/A |
| **(GU)<sub>20</sub>** | 40 | $69 \pm 3.2$ | $1.11 \pm 0.05$ | $528 \pm 28$ | $0.79 \pm 0.03$ | 7.7 |
| **4nt GU loop** | 40 | $1731 \pm 13$ | $2.44 \pm 0.04$ | N/A | N/A | N/A |
| **4 nt UUCG loop** | 40 | $2309 \pm 93$ | $2.52 \pm 0.24$ | N/A | N/A | N/A |
| **8 nt GU loop** | 40 | $1445 \pm 26$ | $2.63 \pm 0.11$ | N/A | N/A | N/A |
| **(GU)<sub>11</sub>** | 22 | $695 \pm 11$ | $1.81 \pm 0.08$ | $833 \pm 8$ | $2.45 \pm 0.05$ | 1.2 |
| **(GU)<sub>12</sub>** | 24 | $359 \pm 7$ | $1.78 \pm 0.06$ | $773 \pm 25$ | $2.4 \pm 0.05$ | 2.2 |
| **(GU)<sub>30</sub>** | 60 | $13 \pm 1$ | $0.87 \pm 0.06$ | $680 \pm 56$ | $0.54 \pm 0.03$ | 52.3 |
| **(AC)<sub>20</sub>** | 40 | $3275 \pm 87$ | $1.51 \pm 0.07$ | N/A | N/A | N/A |
| **(AC)<sub>30</sub>** | 60 | $3603 \pm 93$ | $1.67 \pm 0.08$ | N/A | N/A | N/A |
| **3' UTR** | 321 | $69 \pm 4$ | $0.89 \pm 0.04$ | N/A | N/A | N/A |
| **3' UTR GU mut** | 321 | $141 \pm 5$ | $1 \pm 0.03$ | N/A | N/A | N/A |
| **3' UTR GU stretch mut** | 321 | $204 \pm 12$ | $1.1 \pm 0.07$ | N/A | N/A | N/A |
| **hemi-methylated DNA substrate** | 40 | $149 \pm 2$ | $2.1 \pm 0.05$ | N/A | N/A | N/A |

$K_d^{app}$ values are mean ± Standard Error of the Mean (SEM) (n=3). N/A, not applicable.

**FIGURE 2.**
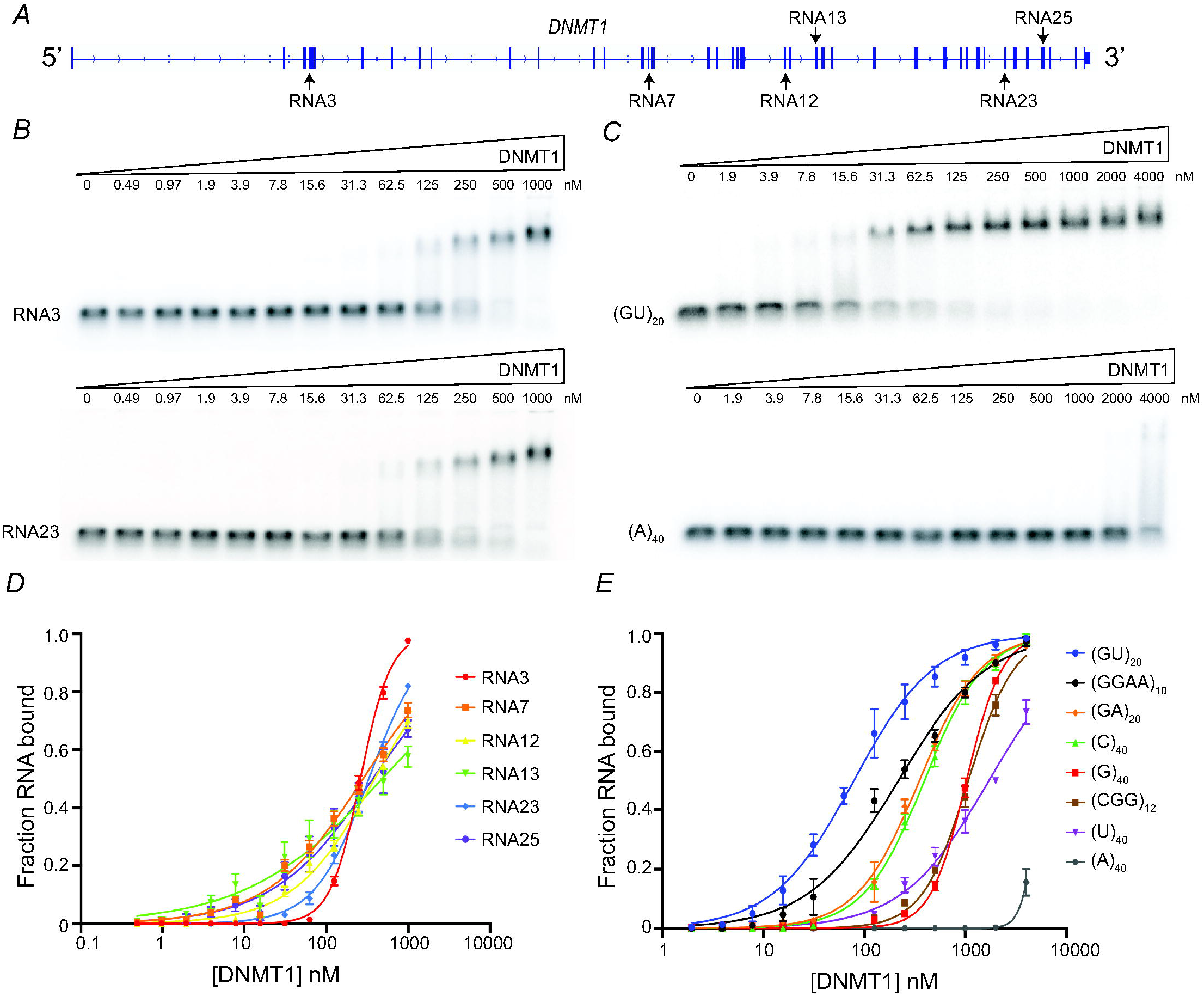
DNMT1 binds to a variety of RNAs with different affinities in vitro. (*A*) Schematic of the location of in vitro transcribed RNAs. (*B*) Representative EMSAs with radiolabeled trace amounts of RNA3 and RNA23 with increasing amounts of DNMT1 protein. (*C*) Representative EMSAs with radiolabeled trace amounts of (GU)_20_ and A_40_ with increasing concentrations of DNMT1. (*D*) Binding curves for all *DNMT1* mRNA constructs. (*E*) Binding curves for all synthetic 40-mer RNA oligonucleotides. In D and E, points represent mean values and error bars represent SD for n = 3 replicates.

Given that the natural RNA sequences tested bound DNMT1 with similar affinities, we used a series of synthetic 40-mer RNAs with the aim of identifying any sequence specificity. Indeed, our binding assays revealed substantial specificity. Among the homopolymers, DNMT1 had the highest affinity for (C)_40_ and undetectable binding to (A)_40_ (**Fig 2C, E, Table 1**). The G4 RNA (GGAA)_10_ bound with somewhat higher affinity than (GA)_20_, which has the same base composition but cannot form G4s. The observed higher affinity for G4s is consistent with previous studies showing direct interactions of DNMT1 with both RNA and DNA G4s (Mao et al. 2018; Zhang et al. 2015). Additionally, we found that DNMT1 bound to many of these RNAs with equal or higher affinity than to a hemimethylated DNA substrate of equal length (**Table 1**). Interestingly, DNMT1 showed particularly high affinity to (GU)_20_ (69 nM *K*_*d*_^*app*^) (**Fig 2C, E, Table 1**). Intrigued by this result, we further investigated the nature of DNMT1’s affinity for GU repeats.

### DNMT1 interacts with RNAs capable of forming non-canonical G4s

Since G and U can form wobble base pairs, it seemed possible that DNMT1 might be recognizing a stem-loop structure with G-U pairs in the stem and a hairpin loop comprised of GU repeats. We, therefore, designed hairpins with different sequence compositions and loop sizes, and we added GU repeats within the loops to tease apart potential binding preference of the loop vs stem (**Supplemental Fig S4A, Supplemental Table 1**). All stem-loop constructs ran with much higher mobility on a native gel compared to (GU)_20_, indicating that (GU)_20_ does not in fact form a stem-loop structure under these conditions (**Supplemental Fig S4B**). This idea was further supported by the fact that while DNMT1 bound to all these stem-loop structures, it had very low affinity for all constructs tested, K_d_^app^ > 1 μM (**Supplemental Fig S4C, Table 1**). These results were surprising given previous reports that DNMT1 preferentially bound RNA stem-loop structures (Di Ruscio et al. 2013).

Recently, another study demonstrated that GU repeats can fold into a structure termed a pUG-fold (Roschdi et al. 2021). pUG-folds are non-canonical parallel G4 structures with a left-handed backbone in which U’s are looped out to allow the formation of three stacked G-quartets capped by a terminal U-quartet. As with canonical G4s, the pUG-fold requires K^+^ ions in order to fold stably (**Fig 3A**). To test for a pUG-fold structure in (GU)_20_, we first ran native gels in K^+^ vs Li^+^ to determine if its folding was K^+^-dependent. Indeed, (GU)_20_ ran with much faster mobility in K^+^ compared to Li^+^, indicating it is forming a K^+^-dependent compact structure (**Fig 3B**). Furthermore, circular dichroism of (GU)_20_ gave a distinct G4-like spectrum in K^+^ but not in Li^+^ (**Fig 3C**). Together, these experiments demonstrate that (GU)_20_ is capable of forming a pUG-fold.

**FIGURE 3.**
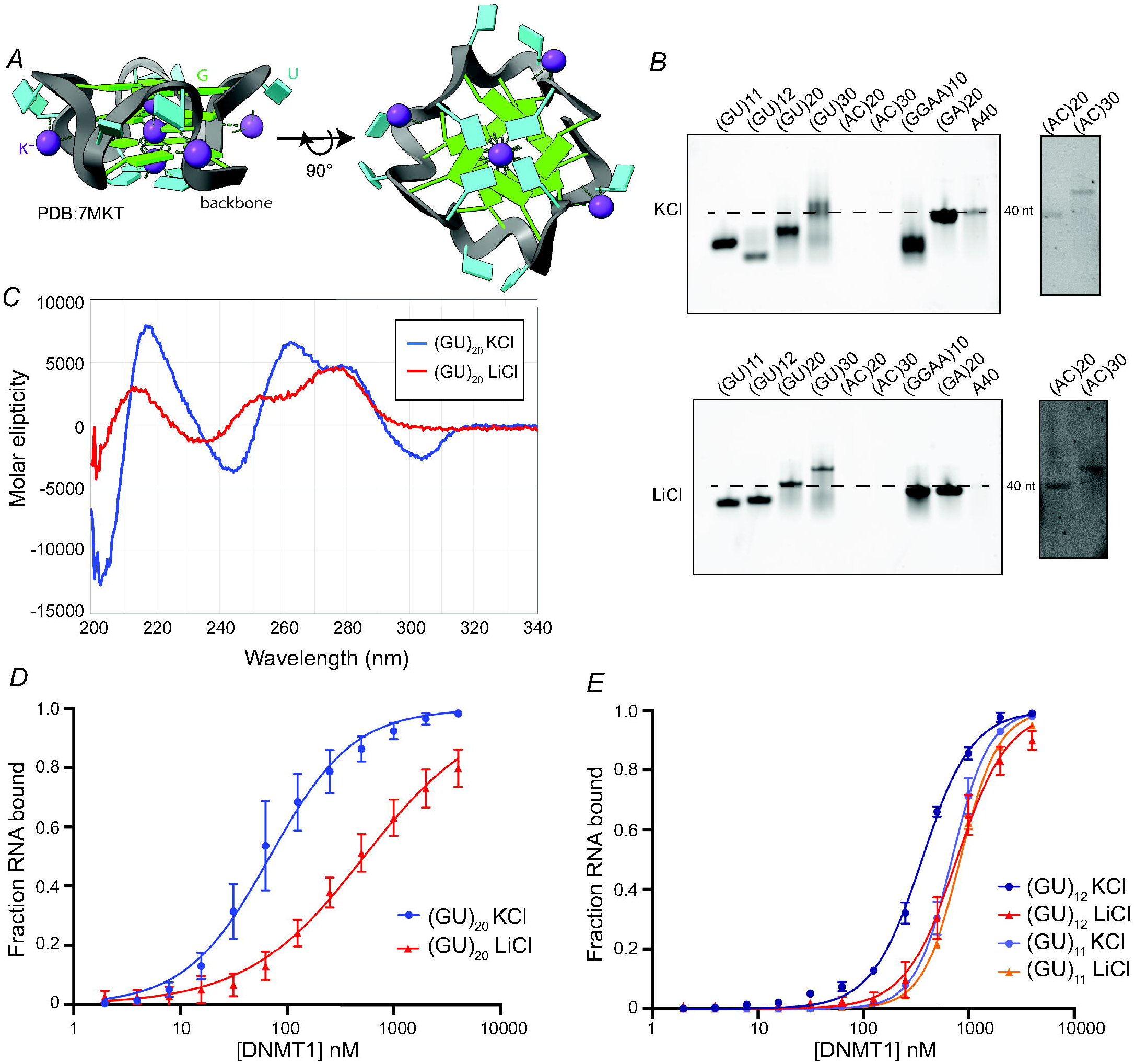
DNMT1 binds with high affinity to pUG-fold RNA. (*A*) Schematic of a pUG-fold. G and U residues are labeled in green and cyan, respectively. The phosphate backbone is highlighted in grey and coordinated K^+^ ions are indicated by purple spheres. PDB: 7MKT (*B*) Native gel of indicated RNAs with either 100 mM KCl or LiCl included in the folding buffer, the gel running buffer, and the gel itself. A_40_ was used as a marker for an unstructured single-stranded RNA (dashed line). Longer exposure of (AC)_20_ and (AC)_30_ staining is shown to the right, due to the low staining of AC sequences by SybrGold. (*C*) Circular dichroism spectra of (GU)_20_ in 100 mM KCl (blue) and LiCl (red). (*D*) Binding curves of (GU)_20_ bound to DNMT1 in 100 mM KCl (blue) and LiCl (red). (*E*) Binding curves of (GU)_12_ and (GU)_11_ bound to DNMT1 in 100 mM KCl (blue and light purple, respectively) and LiCl (red and orange, respectively). In *D* and *E*, points represent mean values and error bars represent SD for n = 3 replicates.

Next, we tested the affinity of DNMT1 for (GU)_20_ in K^+^ vs Li^+^ to determine if the pUG-fold was necessary for DNMT1 binding. Indeed, (GU)_20_ binding affinity was substantially reduced (8-fold) in Li^+^ compared to K^+^ (**Fig 3D, Table 1**). We conclude that DNMT1 binds both the pUG-folded GU repeats (K^+^) and unfolded GU repeats (Li^+^), with substantial preference for the folded form. Intriguingly, DNMT1 seems to exhibit a specific affinity for pUG-folds compared to canonical G4s, as indicated by the increased affinity for (GU)_20_ over RNAs of equal length that can form canonical G4s (3-fold increase over (GGAA)_10_ and a > 10-fold increase over G40 (see **Fig 2B**)).

Additionally, Roschdi et al. found that 12 GU repeats are required to form a single pUG-fold, while 11 repeats are insufficient. We, therefore, tested the binding of DNMT1 to (GU)_12_ and (GU)_11_ in both K^+^ and Li^+^. As predicted by the pUG-fold hypothesis, (GU)_12_ in K^+^ had a significantly higher affinity for DNMT1 than (GU)_12_ in Li^+^ or (GU)_11_ in either K^+^ or Li^+^ (**Fig 3E**). Importantly, the similar binding curve and affinity of DNMT1 for (GU)_11_ in both K^+^ and Li^+^ ensures that it is the pUG-folding capacity and not the presence of Li^+^ ions/absence of K^+^ ions that affect the binding to DNMT1.

### DNMT1 has a higher affinity for multiple pUG-folds

Interestingly, both (GU)_12_ and (GU)_11_ have significantly reduced affinities (> 5-fold) compared to (GU)_20_, indicating that one pUG-fold alone (GU)_12_ does not confer very high affinity for DNMT1 (**Fig 3E**). To determine if additional GU repeats stimulate tighter DNMT1 binding, we performed binding assays with (GU)_30_. As expected from the GU repeat length-dependent affinity observed in the previous experiment (**Fig 3E**), DNMT1 showed > 4-fold tighter binding to (GU)_30_ (K_d_^app^ = 13 nM) compared to (GU)_20_ (**Fig 4A, Supplemental Fig S5A**,**B, Table 1**). Interestingly, (GU)_30_ should be capable of forming two pUG-folds, indicating that the presence of multiple pUG-folds confers higher affinity. To test if this is indeed the case, we repeated the same binding assay in Li^+^ and found a > 50-fold reduction in affinity, arguing that the increased capacity to form pUG-folds indeed contributes greatly to DNMT1 binding affinity (**Fig 4A, Supplemental Fig S5A**,**B, Table 1**).

**FIGURE 4.**
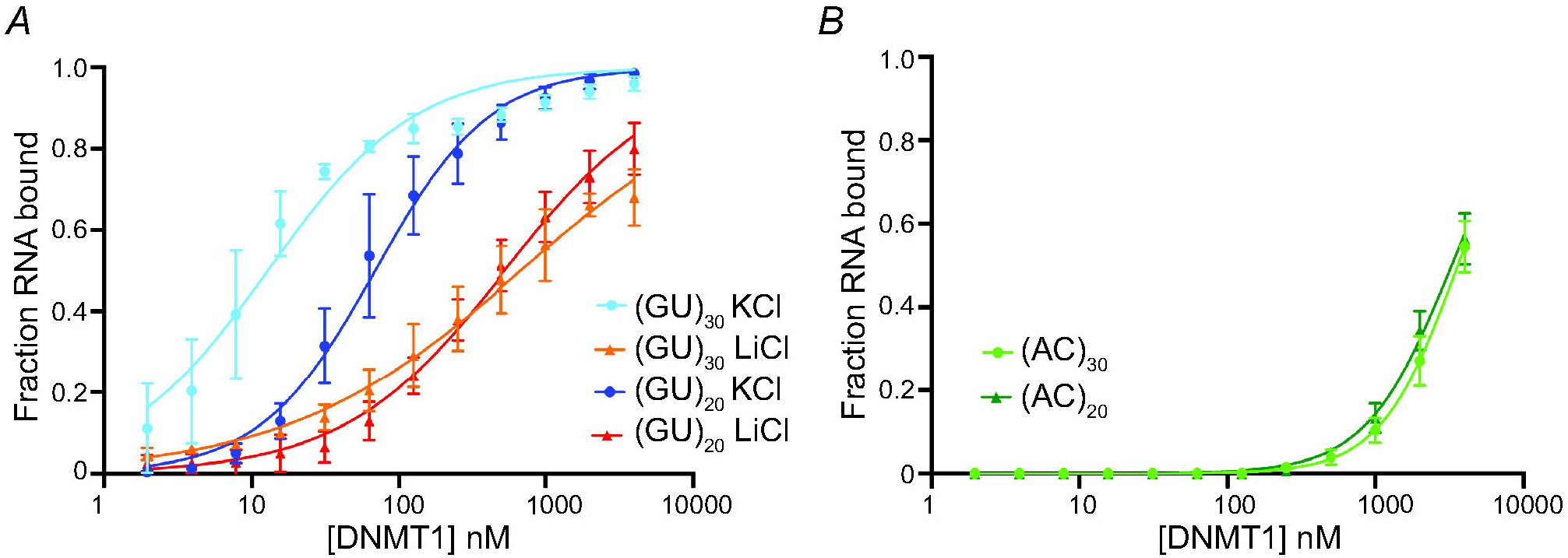
DNMT1 binds to GU repeats in a length-dependent manner. (*A*) Binding curves of (GU)_20_ in KCl and LiCl (blue and red, respectively) and (GU)_30_ in KCl and LiCl (light blue and orange, respectively). Corresponding EMSAs are shown in Supplemental Figure S5. (*B*) Binding curves of (AC)_20_ and (AC)_30_ with DNMT1 (dark green and light green, respectively). Corresponding EMSAs are shown in Supplemental Figure S5. In both panels, points represent mean values and error bars represent SD for n = 3 replicates.

To test if this GU repeat length-specific increase in binding affinity is specific to GU repeats, we performed the same experiment with (AC)_20_ and (AC)_30_. Notably, increased AC repeat content did not increase affinity as seen with increasing GU repeats (**Fig 4B, Supplemental Fig S5C, Table 1**). Together, these results demonstrate that DNMT1 shows a high and specific affinity for pUG-fold RNA.

### GU-repeat containing RNAs inhibit DNMT1 activity

Next, we determined the effect of pUG-fold RNA-binding on DNMT1 enzymatic activity. We first compared the DNMT1 methyltransferase activity on a hemimethylated DNA substrate in the presence of (GU)_20_ or A_40_ in physiologically relevant KCl concentrations (100 mM).

Interestingly, 1 μM (GU)_20_ mostly abolished activity while A_40_ had no effect at the same concentration (**Fig 5A**). We next looked at inhibition by other previously assayed RNAs. The inhibition by (GU)_30_ was stronger than that of (GU)_20_ (**Fig 5B, Supplemental Fig S6**), which is consistent with it having the highest binding affinity of all RNAs tested (**Table 1**). (GU)_12_, which is just long enough to form one pUG-fold, also strongly inhibited DNA methylation (**Fig 5B**). In contrast, (GU)_11_, which is too short to form a pUG-fold, only mildly inhibited activity (**Fig 5B**). These results indicate that pUG-fold RNA is inhibitory to DNMT1 enzymatic activity. Furthermore, a canonical G4 RNA (GGAA)_10_ was also inhibitory, although about 2-fold less than pUG-fold RNA (**Fig 5B**). (GA)_20_, which has the same sequence composition as (GGAA)_10_ but no G4-potential, mildly inhibited activity of DNMT1. Like (A)_40_, (C)_40_ also showed no inhibition of DNMT1 activity, even though it has a much higher affinity for DNMT1 compared to (A)_40_ (**Table 1**). Together, these results argue that G4 RNA, specifically pUG-fold RNA, is particularly inhibitory to DNMT1 enzymatic activity.

**FIGURE 5.**
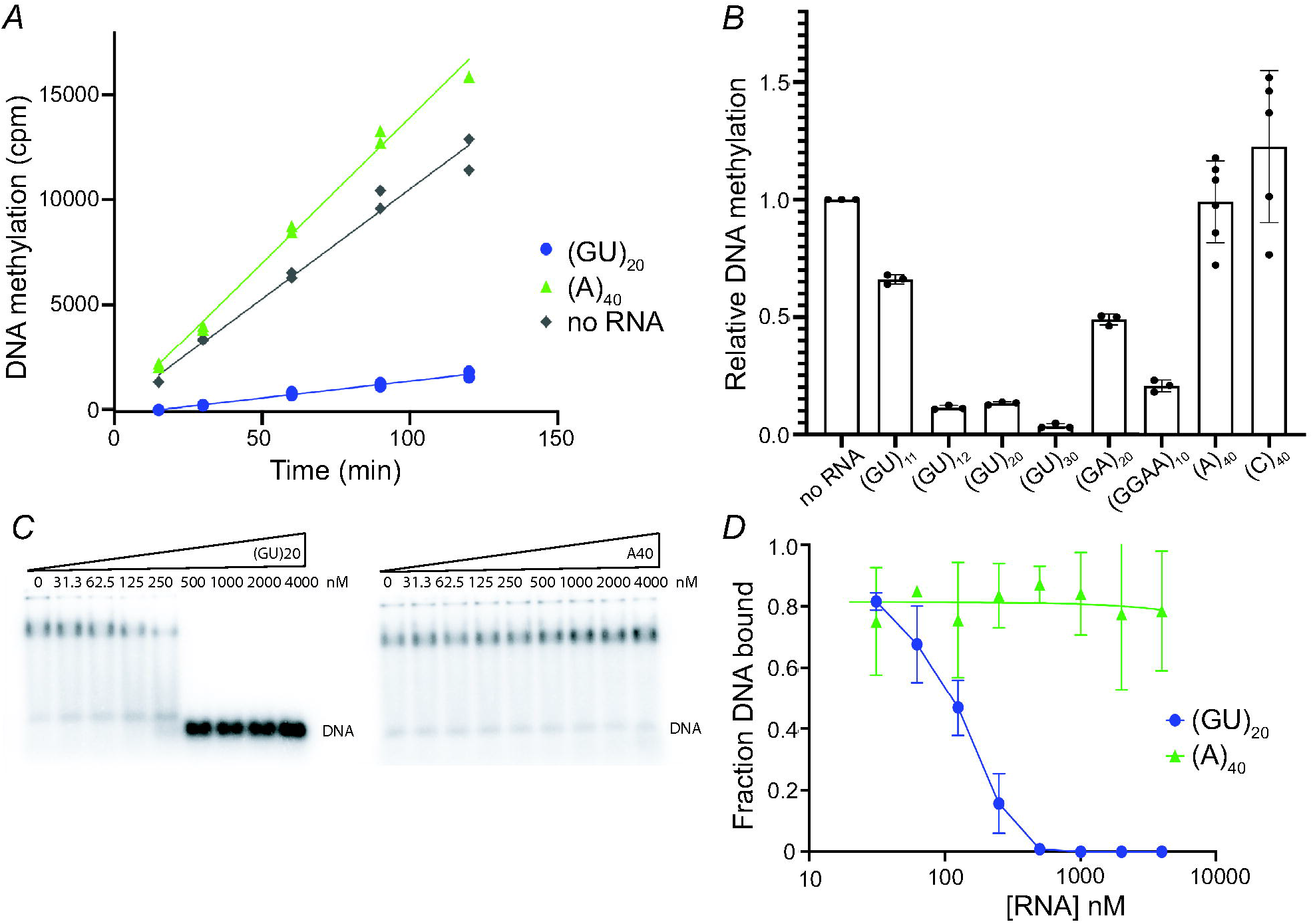
Differential inhibition of DNMT1 methyltransferase activity by different RNAs, with long GU repeats being particularly inhibitory. (*A*) Time course of DNA methylation with 0.25 μM DNMT1, 1 μM hemimethylated DNA substrate, and 1 μM (GU)_20_ or (A)_40_. DNA methylation was measured by incorporation of H^3^SAM and quantified by counts per minute (cpm). Points represent technical replicates. Entire experiment was repeated three times with similar results. (*B*) DNA methylation in the presence of 1 μM of each indicated RNA. Methylation levels have been normalized to amount of methylation measured in the “no RNA” sample. Points represent individual measurements, bars give mean value. Entire experiment was repeated four times with similar results with the exception of (GA)_20_, which showed moderate inhibition in three out of four experiments and strong inhibition in one experiment. (*C*) Representative binding competition EMSAs with trace amounts of hemimethylated DNA substrate and 0.5 μM DNMT1 with increasing concentrations of (GU)_20_ and (A)_40_. (*D*) Binding curves for competition EMSAs shown in *C*, (GU)_20_ (blue) and (A)_40_ (green). Points represent mean values and error bars represent SD for n = 3 replicates. (A)_40_ datapoints are fit with a simple linear regression (green line).

To further understand the inhibition of DNMT1 activity, we measured the ability of (GU)_20_ and A_40_ to compete with the hemimethylated DNA substrate for binding. We saw strong competition for binding by (GU)_20_ with IC_50_ ∼ 100 nM (**Fig 5C, D**). These results indicate that pUG-fold RNA inhibits DNMT1 activity by preventing its hemimethylated DNA substrate from binding. If this inhibition is due to direct competition in the active site or due to an allosteric change remains to be determined.

### DNMT1 binds to GU repeats within its own mRNA

Next, we sought to determine if the affinity of DNMT1 for GU-rich RNAs explains the binding to the *DNMT1* mRNA that we initially observed in the fRIP-seq results. We wondered if the bias in affinity towards the 3’ end of mRNA may be explained by high GU-repeat content. We analyzed the GU-content of the 3’ UTR of the *DNMT1* mRNA and noticed that it is indeed GU-rich with UG/GU being the most frequently occurring dinucleotide (20.8% GU/UG content) (**Supplemental Fig S7**). To test if DNMT1 binds specifically to its own 3’UTR, and if this interaction is GU-repeat dependent, we first performed an EMSA with the full-length in vitro transcribed 321 nt 3’UTR (**Fig 6A**). The 3’UTR had a binding affinity similar to that of (GU)_20_ (both 69 nM *K*_*d*_^*app*^), although the larger size of 3’UTR RNA may contribute to its affinity **(Table 1)**. Mutation of all GU/UG-dinucleotides to AC/CA resulted in a 2-fold reduction in affinity (**Fig 6A**). Furthermore, a more targeted mutation that removed only consecutive GU repeats of three or more gave a larger reduction in affinity (3-fold), indicating that consecutive GU-repeat stretches contribute to binding to DNMT1 (**Fig 6A**). The limited reduction in affinity is not surprising given DNMT1’s general affinity for RNA (e.g., the *DNMT1* mRNA truncation RNA3, which is not very GU-rich).

**FIGURE 6.**
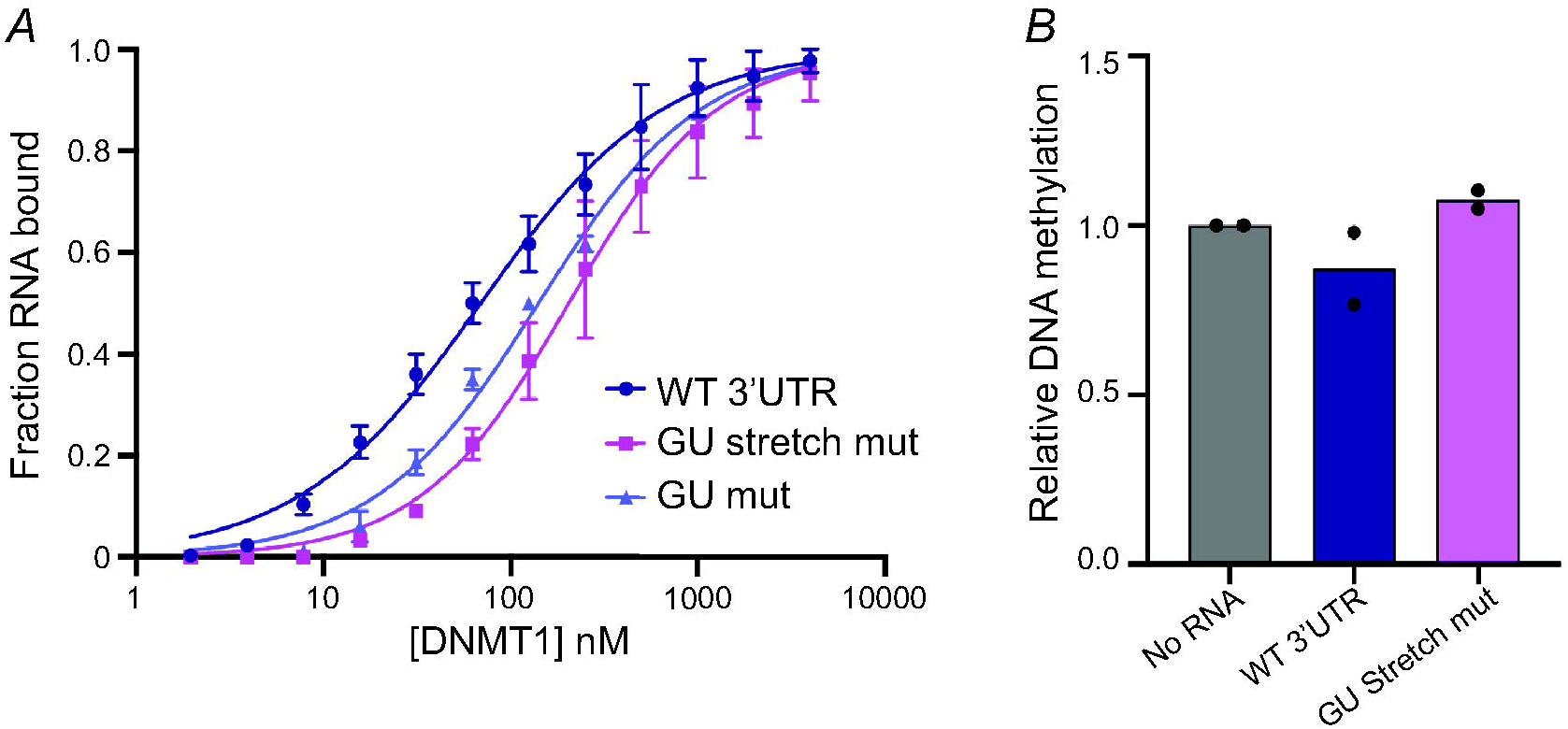
*DNMT1* 3’UTR binds to DNMT1 but does not inhibit activity in vitro. (*A*) Binding curves for the WT 3’UTR (purple) and GU mutant (light purple) and GU stretch mutant (pink). (*B*) Relative DNA methylation in the presence of 1 μM WT 3’UTR (purple) and GU stretch mutant (pink). Methylation levels have been normalized to amount of methylation measured in the “no RNA” (grey) sample. Bars give mean values, with individual data points shown as dots, n=2.

We next tested if the WT 3’UTR or the GU-stretch mutant would influence DNMT1’s methyltransferase activity. Neither RNA affected activity significantly in vitro (**Fig 6B**). The high affinity of the 3’UTR RNA for DNMT1 and its lack of inhibition of activity is in striking contrast to the behavior of pUG-fold RNAs and indicates a different mode of binding.

## DISCUSSION

Here we have uncovered two modes of RNA-binding by DNMT1: a high and specific affinity for pUG-fold RNA and a moderate affinity, seemingly promiscuous binding to many other types of RNA. The pUG-fold mode of binding inhibits DNA methyltransferase activity in an RNA concentration-dependent manner, while the more promiscuous RNA binding has little to no effect. Given the diversity of RNA sequences and structures bound by DNMT1, there may in fact be more than two modes of binding. The second mode of binding is consistent with previous studies showing that DNMT1 binds to multiple different RNAs of varying length and secondary structure with variable effects on DNMT1 enzymatic activity. (Di Ruscio et al. 2013; G Hendrickson et al. 2016; Merry et al. 2015; Zhang et al. 2015) Additionally, we found that many RNAs bind DNMT1 with similar or higher affinity than that of a hemi-methylated DNA substrate (**Table 1**). This preferred affinity for RNA over its natural DNA substrate has been observed previously as well (Di Ruscio et al. 2013).

In addition to DNMT1 having a high affinity for pUG-fold RNA, it seems that RNAs with higher pUG-fold potential, that is, either larger or multiple pUG-folds, confer the highest affinity. To our knowledge, DNMT1 is the first human protein known to bind pUG-folds. It is interesting to note that while DNMT1 has never previously been shown to bind pUG-fold RNA, it has been shown to bind both canonical RNA and DNA G4s. RNA G4s have been shown to inhibit DNMT1 activity by directly competing with the DNA substrate, while DNA G4s have been hypothesized to inhibit DNMT1 through allosteric mechanisms (Mao et al. 2018; Zhang et al. 2015). Furthermore, in our study we note that the strength of inhibition seems correlated with RNA structure rather than with affinity. RNAs with relatively high affinity, e.g., 3 ‘UTR, do not affect DNA methylation. On the other hand, RNAs with similar affinity (compare (C)_40_ and (GU)_12_ (**Table 1**)) have very different effects on activity, with (C)_40_ having no effect and (GU)_12_ being strongly inhibitory.

While non-canonical G4s have been characterized previously, especially in RNA aptamers, there is limited information regarding the affinity of RNA-binding proteins (RBPs) for canonical vs non-canonical G4s (Banco and Ferré-D’Amaré 2021; Vasilyev et al. 2015; Yatime et al. 2015). There is some evidence that the less stable nature of non-G quartets within non-canonical G4s may allow for a more seamless transition between G4 structures and other commonly found RNA secondary structures such as stem-loops (Warner et al. 2014). The increased affinity of DNMT1 for pUG-fold RNA over canonical G4 RNA in our study hints at a potential binding preference for such a hairpin-G4 structure over a G4 alone. Alternatively, it is possible that the increased affinity is driven by the pUG-fold-specific U-propeller loops or the terminally stacked U-quartet. It is also worth noting that natural RNAs with fewer than 12 consecutive UG repeats may still form pUG-folds, because the structure tolerates certain insertions and substitutions (Roschdi et al. 2022), similar to that of previously reported G4 structures (Lightfoot et al. 2019). A comprehensive analysis of how many and what types of insertions/substitutions are allowed is needed.

In this study we found that DNMT1 binds to its own mRNA in two human cell types and that this interaction seems to be specific to the fully spliced transcript in the nucleus. Binding to a spliced transcript was unexpected, because on average DNMT1 fRIP signals for introns were higher than for exons, as expected for a nuclear protein (**FigS2A**). Furthermore, DNMT1 binds with high affinity to its own 3’UTR but does not inhibit methyltransferase activity in vitro. This suggests that the DNMT1-*DNMT1* mRNA interaction could be regulatory in cells, e.g., by influencing the binding of DNMT1 to its chromatin substrate or sequestering mature *DNMT1* mRNA in the nucleus, preventing it from being transported to the cytoplasm for translation. This could represent an autoregulatory mechanism to control DNMT1 levels in the cell.

Recent studies have demonstrated that binding of DNMT1 to a variety of ligands induces a conformational change in the enzyme (Onoda et al. 2022; Ren et al. 2018a, 2021). Additionally, the ATP-dependent methyltransferase DNMT5, a fungal relative of DNMT1, undergoes sequential conformational changes upon binding first to the cofactor SAM, then to ATP and finally to its native DNA substrate (Wang et al. 2022). It is interesting to consider the potential conformational changes that different types of RNA may stimulate when binding to DNMT1. For example, does (GU)_20_ simply occlude the active site when bound to prevent access to its native hemimethylated substrate? Or does it stimulate a conformational change that renders the active site unavailable? To our knowledge, there are no structural studies of DNMT1 bound to RNA. Future structural work on DNMT1/pUG-fold complexes are needed to better understand the nature of this intriguing interaction.

Formation of G4s within the UTRs of the mRNAs of many oncogenes and transcriptional regulators has been linked to translational repression (Huppert et al. 2008; Kumari et al. 2007; Shahid et al. 2010). Furthermore, disruption of RNA G4s within the 5’UTR of multiple oncogenes was shown to increase cancer susceptibility (Kosiol et al. 2021; Wolfe et al. 2014). It is interesting to consider that RNA G4s may play additional regulatory roles in trans through interactions with epigenetic regulators. Like DNMT1, PRC2 – another chromatin-associated protein that promotes formation of heterochromatin – has been shown to bind and be regulated by G4 RNA (Beltran et al. 2019; Kaneko et al. 2014; Long et al. 2017, 2020; Wang et al. 2017). Curiously, the increased affinity of DNMT1 for pUG-fold G4s over canonical G4s is in contrast to the preferred affinity of PRC2 for canonical G4s vs GU-repeat RNA (> 3-fold increased affinity for canonical G4s (GGAA)_10_ vs (GU)_20_) (Wang et al. 2017). It is interesting to speculate that binding of G4s to chromatin-associated proteins may be an evolved mechanism for silencing of gene expression and that the difference in binding preference for different types of G4s could reflect a fine-tuning of RNA-based inhibition of different epigenetic factors.

## MATERIALS AND METHODS

### Cell culture and crosslinking

Ten million K562 cells (ATCC: CCL-243) or iPSCs (WTC-11) were grown for each experiment. Cells were spun down for 5 min at 500 x g and washed with 10 ml cold PBS. Cells were spun down and the supernatant was removed. Cells were fixed in 40 ml PBS + 0.1 % formaldehyde at room temperature for 10 min. The formaldehyde-solution was then quenched with 2 ml 2.5 M Glycine (drops added slowly) followed by rotation at room temperature for 5 minutes. Cells were spun down at 500 x g for 5 min at 4°C. Supernatant was removed and cells were resuspended in 10 ml cold PBS + 1X Protease inhibitor (Pierce tablets). Cells were spun down for 5 min at 500 x g and supernatant was discarded. Resuspend pellet in 10 mL ice-cold PBS + 1X PIC. Cells were spun down at 500 x g for 5 min and pellets were flash frozen in liquid nitrogen and stored at −80**°**C or immediately used.

### fRIP

Crosslinked cell pellets were thawed and resuspended in 1 mL cold fRIP lysis buffer (50 mM Tris pH 8, 150 mM KCl, 0.1% SDS, 1% Triton-X, 5 mM EDTA, 0.5% sodium deoxycholate, 0.5 mM DTT, 1x Protease Inhibitor Cocktail (Roche), 100 U/mL RNasin Plus RNase inhibitor (Promega)). Samples were rotated at 4°C for 10 minutes and then sonicated at 4°C in a Bioruptor UCD-200 (diagenode) on the high setting using a 30 seconds on/30 seconds off cycle for 15 minutes. Cells were spun down at 13,000 RPM for 10 minutes at 4°C. Pellet was discarded and 1 ml of cold fRIP wash buffer (25 mM Tris pH 7.5, 150 mM KCl, 5 mM EDTA, 0.5% NP-40, 0.5 mM DTT, 1x Protease Inhibitor Cocktail (Roche), 100 U/mL RNasin Plus RNase inhibitor (Promega)) was added to 1 ml of supernatant to bring volume to 2 ml. Sample was filtered through a 0.45 micron filter into a new 2-mL tube and sample was split evenly between two new tubes (each with ∼5 million cells worth of lysate).

Pierce Protein A/G dynabeads (Thermo Scientific) were equilibrated with cold fRIP wash buffer (25 mM Tris pH 7.5, 150 mM KCl, 5 mM EDTA, 0.5% NP-40, 0.5 mM DTT, 1x Protease Inhibitor Cocktail (Roche), 100 U/mL RNasin Plus RNase inhibitor (Promega)) and 25 ul beads were added to each tube of lysate. Tubes were rotated at 4 °C for 30 minutes. Beads were pulled down with a magnetic rack and supernatant was transferred to new tube (Precleared lysate). Two 25 ul aliquots of precleared lysate were taken here as input and western blot samples. Precleared lysates were flash-frozen in liquid nitrogen and stored at −80°C until further use or used directly for IP.

One tube of pre-cleared lysate was thawed for each IP condition. 6 ug of appropriate antibody was added to the lysate (DNMT1: Abcam 19905 Lot No: GR3271225, Abcam 13537 Lot No: GR3184365, HNRNPH: Bethyl A300-511A, Lot: 2). Tubes were vortexed gently and then rotated at 4 °C for 2 hours. 50 μl of equilibrated protein A/G magnetic beads were added to each tube of lysate. Samples were incubated for 1 h at 4 °C while rotating. Beads were pulled and supernatant discarded. Beads were washed 4 times in 1 mL fRIP wash buffer (25 mM Tris pH 7.5, 150 mM KCl, 5 mM EDTA, 0.5% NP-40, 0.5 mM DTT, 1x Protease Inhibitor Cocktail (Roche), 100 U/mL RNasin Plus RNase inhibitor (Promega)), each time rotating at 4 °C for 10 minutes. Before removing last wash, 100 μl of beads were transferred to a new tube. This 10 % aliquot served as an IP sample for a western blot, and the remaining 90 % was used for isolating RNA. Beads were pulled down for both the 900-μL and 100-μL samples and supernatant removed. Beads were either stored at −20 °C or continued with for reverse crosslinking. The input and IP samples were thawed and 31 μl water was added to the input and 56 μl water was used to resuspend the IP beads. To each sample, 33 μl 3X RCL Buffer (3X PBS, 6 % N-lauroyl sarcosine, 30 mM EDTA, 15 mM DTT), 0.2 mg Proteinase K (Invitrogen) and 1 μl RNasin Plus RNase inhibitor (Promega) was added. Samples were incubated at 42 °C for 1 h and then 55 °C for 1 h. RNA was isolated via Phenol/Chloroform extraction followed by ethanol precipitation. Pellet was resuspended in 15 μl nuclease-free water.

### Library construction and next generation sequencing

Purified RNA from input and IP samples were reverse transcribed, PCR amplified and barcoded using the KAPA RNA HyperPrep kit with RiboErase. Resulting cDNA libraries were paired-end sequenced on a Illumina NextSeq 500 at a depth of 20-40 million reads/sample. All whole cell fRIP experiments had 3 replicates and the nuclear fRIP had 4 replicates.

### fRIP-seq computational analysis

Read mapping and QC of sequenced samples was handled by nf-core/chipseq v1.1.0 pipeline (Ewels et al. 2020). Reads were mapped to hg38 (Gencode v32 annotation) using bwa and the default nf-core parameters. For gene level enrichment analysis, counts were quantified using Rsubread featureCounts and differential enrichment was calculated with DESeq2. All code to reproduce this analysis is available in the git repository: https://github.com/ljansson/dnmt1.git.

### Nuclear fractionation

Protocol was adapted from the Bönisch Lab (Arrigoni et al. 2016). Twenty million K562 cells were grown/experiment. Cells were crosslinked and pelleted as described for the whole cell fRIP above. Crosslinked pellets were thawed and gently resuspend in 1 mL cold Farnham Lab (FL) buffer (5mM PIPES pH 8.0, 85 mM KCl, 0.5% NP-40, 1X Protease Inhibitor Cocktail (Roche)). Cells were then sonicated in a Covaris Ultrasonicator for 30 s with 75W peak power, 2% duty factor and 200 cycles/burst at 4-10 °C.

Cells were spun down at 1000 x g for 5 min at 4 °C. Supernatant was removed and nuclear pellet was washed once with 1 ml 1X FL buffer (5mM PIPES pH 8.0, 85 mM KCl, 0.5% NP-40, 1X Protease Inhibitor Cocktail (Roche)). Pellet was spun down at 1000 x g for 5 min at 4 °C. Supernatant was removed and pellet was resuspended in 500 ul of 1X fRIP lysis buffer (50 mM Tris pH 8, 150 mM KCl, 0.1% SDS, 1% Triton-X, 5 mM EDTA, 0.5% sodium deoxycholate, 0.5 mM DTT, 1x Protease Inhibitor Cocktail (Roche), 100 U/mL RNasin Plus RNase inhibitor (Promega)). fRIP was then performed as described above for whole-cell pellets. Nuclear fractionation was confirmed by western blot. Antibodies used for western blot include: DNMT1 (Abcam 19905), EZH2 (Cell Signaling Technologies 5246S) and β-actin (Sigma-Aldrich A5441).

### Full-length human MBP-DNMT1 expression

Sf9 cells were used to generate baculovirus stocks using the Bac-to-Bac system (Life Technologies), according to the manufacturer’s instructions. A baculovirus stock carrying the gene for DNMT1 was used to infect 4–12 l of Sf9 cells at a density of 2 million cells/ml in Sf-900™ III SFM (Invitrogen cat # 12658–027). Multiple 1 l cell cultures in 4 l flasks (VWR cat # 32645–044) were incubated 72 h at 27 °C, 130 revolutions per minute (rpm). Cells were harvested by centrifugation for 20 min at 2000 relative centrifugal force (RCF) (JLA-8.1, Beckman) at 4°C, frozen in liquid nitrogen and stored at −80 °C until protein purification.

### Full-length human MBP-DNMT1 purification

DNMT1 cell pellet was weighed and 50 ml of lysis buffer (10 mM Tris-HCl pH 7.5, 250 mM NaCl, 0.5% NP40, 1 mM TCEP) was added to every 6 g of pellet. Pellet was gently resuspended followed by incubation at 4 °C while rotating for 15 min. Cells were spun down at 18000 rpm for 30 min at 4 °C. Pellet was discarded and supernatant was incubated with equilibrated amylose resin (NEB, E8021L) (1 ml resin/ 2 g pellet) for 90 min at 4 °C while rotating. Sample was then washed in batch with first 10 cv lysis buffer, 16 cv high salt buffer (10 mM Tris-HCl pH 7.5, 500 mM NaCl, 1 mM TCEP), 16 cv low salt buffer (10 mM Tris-HCl pH 7.5, 150 mM NaCl, 1mM TCEP) with 5 min spins at 1000 x g between each wash. MBP-DNMT1 was eluted in low salt buffer + 10 mM maltose. Elution was concentrated and cleaved overnight with 1:50 w/w ratio PreScission protease at 4 °C. Following cleavage, DNMT1 was purified by ion-exchange chromatography on a HiTrap 5 ml Heparin HP column (Cytiva) and fractions were pooled and concentrated. Heparin-purified sample was then further purified by size-exclusion chromatography on a Sepharose 6 increase 10/300 GL column (Cytiva). Fractions were pooled and concentrated and aliquots of pure DNMT1 were flash frozen and stored at −80 °C until use.

### In vitro transcription

PCR primers were designed to PCR amplify *DNMT1* mRNA fragments from a full-length human DNMT1 cDNA plasmid (plasmid details available upon request). PCR products were purified using the E.Z.N.A. Cycle Pure kit (Omega bio-tek) and used as templates for T7 in vitro transcription. The in vitro transcription reaction was incubated overnight at room temp (37 °C for 3’ UTR constructs). The reaction then incubated with Turbo DNase for 15 min. The RNA was then isolated using phenol/chloroform extraction and ethanol precipitation followed by PAGE purification.

### Electrophoretic mobility shift assay

In vitro transcribed RNAs were CIP-treated before end-labeling. 50 pmol of in vitro transcribed or synthetic RNA was incubated with 10 units T4 PNK (NEB) and 2 μl γ-^32^P-ATP in 1X PNK buffer and incubated for 1 h at 37 °C. Radiolabeled RNAs were purified using QuickSpin columns (Roche) and radioactivity was measured by scintillation counter. End-labeled RNA was folded by first heating to 95C for 5 min followed by snap cooling on ice for 2 min and then incubated for 30 min at 37 °C in 1X refold buffer (50 mM Tris-HCl pH 7.5, 100 mM KCl, 2.5 mM MgCl_2_, 0.1 mM ZnCl_2_, 2.0 mM BME, 0.05 % NP40, 5 % v/v glycerol). Each binding reaction contained the indicated amount of DNMT1 plus 1000 cpm of end-labeled RNA in binding buffer (50 mM Tris-HCl pH 7.5, 100 mM KCl, 2.5 mM MgCl_2_, 0.1 mM ZnCl_2_, 2.0 mM BME, 0.05 % NP40, 5 % v/v glycerol, 0.1 mg/mL BSA (NEB), 0.1 mg/mL Baker’s Yeast tRNA (Sigma R5636)). Reactions were incubated for 1 hour at 30 °C. Samples were run on a 1 % agarose gel in 1X TBE and run for 90 min at 4 °C at 66 V. The gel was dried and exposed overnight followed by imaging on a Typhoon FLA 9500 scanner (GE). All EMSAs were performed in triplicate. EMSAs were quantified using either ImageQuant or GelAnalyzer 19.1 and binding curves were generated using Prism 9. 40-mer RNAs were ordered from Dharmacon.

### Binding competition assays

RNAs were folded by first heating to 95 °C for 5 min followed by snap cooling on ice and then incubated for 30 min at 37 °C in 1X refold buffer (50 mM Tris-HCl pH 7.5, 100 mM KCl, 2.5 mM MgCl_2_, 0.1 mM ZnCl_2_, 2.0 mM BME, 0.05 % NP40, 5 % v/v glycerol) Each binding reaction contained 0.5 μM DNMT1, 1000 cpm of end-labeled DNA and the indicated concentration of cold RNA in 1X binding buffer (50 mM Tris-HCl pH 7.5, 100 mM KCl, 2.5 mM MgCl_2_, 0.1 mM ZnCl_2_, 2.0 mM BME, 0.05 % NP40, 5 % v/v glycerol, 0.1 mg/mL BSA (NEB), 0.1 mg/mL Baker’s Yeast tRNA (Sigma R5636)). DNA was directly diluted in 1X folding buffer before incubating with DNMT1 and cold RNA. Hemi-methylated DNA was ordered pre-annealed from IDT: ((5’TA(5mC)GTATC(5mC)GTATC(5mC)GGTTA(5mC)GTATC(5mC)GAATC(5mC)GTAC(5 mC)GT 3’ / 5’ ACGGTACGGATTCGGATACGTAACCGGATACGGATACGTA 3’)). CpG-flanking sequences were designed to promote ideal binding/activity by DNMT1 (Adam et al. 2020).

Reactions were incubated for 1 hour at 30°C. Samples were run on a 1 % agarose gel in 1X TBE and run for 90 min at 4°C at 66V. The gel was dried and exposed overnight followed by imaging on a Typhoon FLA 9500 scanner (GE). All competition EMSAs were performed in triplicate. Binding competition assays were quantified using GelAnalyzer 19.1 and binding curves were generated using Prism 9.

### In vitro methylation assay

Single point biochemistry assays were performed by incubating 0.25 μM DNMT1, 1.0 μM hemimethylated DNA (same as used for binding competition EMSAs), 1.0 μM folded RNA in 1X binding buffer (50 mM Tris-HCl pH 7.5, 100 mM KCl, 2.5 mM MgCl_2_, 0.1 mM ZnCl_2_, 2.0 mM BME, 0.05% NP40, 5% v/v glycerol, 0.1mg/mL BSA (NEB), 0.1 mg/mL Baker’s Yeast tRNA (Sigma R5636)) and 1.0 μM [^3^H] SAM (82.3 Ci/mmol, PerkinElmer). First, buffer, DNA, [^3^H] SAM were combined in a master mix and aliquoted to appropriate tubes, followed by addition of indicated RNA. Reaction was then started by addition of protein. This order-of-addition was designed to ensure that the SAM, which is dissolved in acid, was neutralized with buffer before the protein was added. Reactions were incubated for 2.5 hours at 37 °C. Reactions were cooled to 4°C for 30 seconds and quenched with 1.2 mM unlabeled SAM (NEB) and blotted onto Hybond-XL membrane. Membranes were allowed to air dry, then were batch washed (8 mL wash buffer per membrane) 3x 50 mM KH_2_PO_4_, 1x 80 % ethanol, 1x 100 % ethanol. Membranes were allowed to air dry, then were soaked in ScintiSafe Econo 1 scintillation cocktail. Beckman LS 6500 scintillation counter was used to measure ^3^H incorporation.

Inhibition pattern of various RNAs were determined by incubating 0.25 μM DNMT1, 1 μM hemimethylated DNA, 1 μM [^3^H] SAM (82.3 Ci/mmol, PerkinElmer) and various amounts of refolded RNA in 1X binding buffer. The same incubation, quenching, blotting, and washing protocols were followed as described above. Plots were generated using Prism 9.

### Circular dichroism

RNAs at 0.2 mg/ml were heated at 95 °C for 5 min and then snap cooled followed by folding in 1X refold buffer containing either KCl or LiCl (50 mM Tris-HCl pH 7.5, 100 mM KCl/LiCl, 2.5 mM MgCl_2_, 0.1 mM ZnCl_2_, 2.0 mM BME, 0.05 % NP40, 5 % v/v glycerol) at 37°C for 30 min. 150 μl of sample was added to a 0.5 mm pathlength cuvette and loaded into a ChirascanPlus Circular Dichroism and Fluorescence Spectrometer (Applied Photophysics) and the CD spectrum was recorded. Either KCl buffer or LiCl buffer alone was used as a baseline control for each salt condition. Measurements were taken from 200 – 340 nm. Cuvettes were rinsed with 3X buffer between each sample. When switching between KCl buffer and LiCl buffer, cuvette was additionally washed 2X with H_2_O and once with 100 % ethanol. Baselines were subtracted from experiment spectra and circular dichroism was converted to molar ellipticity.

### Native gel electrophoresis

RNAs were heated at 95 °C for 5 min and then snap cooled followed by folding in 1X refold buffer containing either KCl or LiCl (50 mM Tris-HCl pH 7.5, 100 mM KCl/LiCl, 2.5 mM MgCl_2_, 0.1 mM ZnCl_2_, 2.0 mM BME, 0.05 % NP40, 5 % v/v glycerol) at 37 °C for 30 min. RNAs were loaded onto a 10 % Native PAGE gel containing either 100 mM KCl or LiCl in 0.5 X TBE. Samples were run at 80 V for 1 h in 0.5X TBE buffer plus 100 mM KCl/LiCl at room temperature. Gels were post-stained with SybrGold for 15 min and imaged by a FluorChem R imager (proteinsimple).

Folding of DNMT1 mRNA in vitro transcription products were analyzed by loading onto a 6 % PAGE gel without additional salt in gel and loading buffer.

## Acknowledgements

We thank core facility directors A. Scott (BioFrontiers Next Generation Sequencing Facility), T. Nahreini (Biochemistry Cell Culture Facility and Flow Cytometry Shared Core) and A. Erbse (Shared Instruments Pool in Biochemistry). We thank Sam Butcher and members of his lab for helpful conversations. We also thank J.P. Ouyang from the Parker lab and members of the Cech Lab for thoughtful discussions. This work was supported by NIH grant (5K00CA212439-06) to LJF, NIH T32 training grant (GM142607) to JJS and NIH NIGMS (R00GM132544) to VK. MJS is a member of the Interdisciplinary Quantitative Biology PhD program. JLR is a Howard Hughes Medical Institute (HHMI) Faculty Scholar and TRC is an investigator of the HHMI.

## Author Contributions

LJF, CIS and TRC conceived the study and designed experiments. LJF, CIS and JJS performed experiments and analyzed data. LJF and MJS analyzed fRIP-seq results. ARG performed DNMT1 SF9 cell culture and protein expression. LJF and TRC wrote the manuscript with input from all authors.

